# Preserving Full Spectrum Information in Imaging Mass Spectrometry Data Reduction

**DOI:** 10.1101/2024.09.30.614425

**Authors:** Roger A.R. Moens, Lukasz G. Migas, Jacqueline M. Van Ardenne, Eric P. Skaar, Jeffrey M. Spraggins, Raf Van de Plas

## Abstract

**Motivation:** Imaging mass spectrometry (IMS) has become an important tool for molecular characterization of biological tissue. However, IMS experiments tend to yield large datasets, routinely recording over 200,000 ion intensity values per mass spectrum and more than 100,000 pixels, *i*.*e*., spectra, per dataset. Traditionally, IMS data size challenges have been addressed by feature selection or extraction, such as by peak picking and peak integration. Selective data reduction techniques such as peak picking only retain certain parts of a mass spectrum, and often these describe only medium-to-high-abundance species. Since lower-intensity peaks and, for example, near-isobar species are sometimes missed, selective methods can potentially bias downstream analysis towards a subset of species in the data rather than considering all species measured.

**Results:** We present an alternative to selective data reduction of IMS data that achieves similar data size reduction while better conserving the ion intensity profiles across all recorded *m/z* -bins, thereby preserving full spectrum information. Our method utilizes a low-rank matrix completion model combined with a randomized sparse-format-aware algorithm to approximate IMS datasets. This representation offers reduced dimensionality and a data footprint comparable to peak picking, but also retains complete spectral profiles, enabling comprehensive analysis and compression. We demonstrate improved preservation of lower signal-to-noise-ratio signals and near-isobars, mitigation of selection bias, and reduced information loss compared to current state-of-the art data reduction methods in IMS.

## Introduction

Imaging mass spectrometry (IMS) is an analytical imaging technology that enables molecular mapping of complex biological samples, such as tissues, biofilms, or dispersed cells [Caprioli et al., 1997, McDonnell and Heeren, 2007, Esselman et al., 2023, Perry et al., 2022, Bien et al., 2022]. IMS combines the sensitivity and specificity of mass spectrometry with spatial information. It enables researchers to concurrently measure the distribution of hundreds to thousands of molecular species throughout tissue sections or other heterogeneous samples without the need for labeling target molecules [Caprioli et al., 1997, McDonnell and Heeren, 2007, Buchberger et al., 2018, Aichler and Walch, 2015]. This capability holds strong potential for probing the lipidomic, glycomic, metabolomic, and proteomic content of biological samples across a wide range of applications, spanning from fundamental research in biology and medicine to the development of novel diagnostics and therapeutics [Rubakhin et al., 2005, Vaysse et al., 2017, Kaspar et al., 2011].

However, the outstanding multiplexing capability of IMS-capable instruments generates vast amounts of data, often containing spatially resolved information for thousands of molecular species in a single experiment [Caprioli et al., 1997, Spraggins et al., 2019, Alexandrov, 2012]. The volume and high-dimensionality of IMS data present significant challenges in data processing, analysis, and interpretation [Alexandrov, 2012]. One of the primary challenges is data reduction, as raw IMS datasets typically consist of hundreds of thousands to millions of spatial locations, *i*.*e*., pixels, each associated with a mass spectrum containing hundreds of thousands of ion intensities. These datasets contain a mixture of high- and low-intensity peaks as well as features with varying signal-to-noise ratios. Managing such large datasets requires effective data reduction techniques that extract meaningful information while minimizing computational burden and storage demands [Verbeeck et al., 2020].

Current data reduction approaches can be broadly classified into acquisition-time and post-acquisition data reduction methods. Acquisition-time data reduction involves reducing data during acquisition, typically resulting in a selective representation rather than describing full spectra. For instance, time-of-flight (TOF) mass spectrometers may retain only ion intensity values above a certain threshold, discarding other measured *m/z* -bins. These representations, while reduced, are often not well-suited for downstream analysis due to their still relative-large size, unstructured feature selection, and sparse matrix format, commonly requiring reconstruction to the full mass domain (intensity values for all *m/z* -bins) and subsequent data re-reduction. Moreover, this process is usually “hard-coded” into the instrument and not under user control.

Post-acquisition data reduction is user-controlled and includes techniques such as peak picking (*i*.*e*., selecting a subset of features or masses), spectral integration (*i*.*e*., combining features over small mass ranges), and spatial cropping (*i*.*e*., selecting a spatial subset of interesting pixels and spectra) [Alexandrov, 2012, Anderson et al., 2016, Monchamp et al., 2007]. Note that ion intensity integration sometimes already happens at the detector and/or instrumental level, for example, to counteract space-charge effects. These post-acquisition methods usually aim to convert full spectrum IMS data, that is IMS data with ion intensity values for each *m/z* -bin in the measured mass range, into a more manageable representation reporting only the signal features deemed important, often certain peaks, and with a reduced memory footprint such that it is practical for subsequent analysis [Verbeeck et al., 2020, Alexandrov, 2012]. However, methods like peak picking can miss or inadequately capture certain peaks in the full spectra. These missed signals often include low-intensity or low signal-to-noise-ratio (*SNR*) peaks and near-isobars (*i*.*e*., peaks with nearly identical *m/z* but representing different molecular species, and sometimes presenting as ‘shoulders’ in a peak profile). This is often due to their focus on the abundance of signals rather than considering the structured signal presence or absence across measurements. Furthermore, their selective nature can introduce bias by limiting downstream analysis to a subset of (often more abundant) molecular species rather than considering all measured species. Although recent efforts have been made to improve peak-picking accuracy and robustness [González-Fernández et al., 2023], challenges persist, particularly in handling near-isobaric species and low-intensity peaks.

Here, we introduce a novel data model for IMS measurements that addresses missing values through sparse-format-aware low-rank matrix approximation. This approach offers an alternative to traditional data reduction methods for IMS, mitigating selection bias and minimizing information loss early in the analysis pipeline by preserving full spectrum information rather than only selective sub-windows of the measured mass range. Additionally, to handle the computational demands and large memory requirements of modern IMS datasets, we explore the use of randomization strategies to optimize low-rank factorization methods and implement obtaining such a representation of an IMS dataset.

## Availability

The source code for the methods introduced in this work is available at https://github.com/vandeplaslab/full_profile, and the data is available on request. We provide several sparsity-format-aware implementations of the Singular Value Thresholding (SVT) and Fixed Point Continuation (FPC) algorithms, developed in object-oriented Python for both CPU and GPU. These algorithms have been optimized for efficiency given available computational resources.

## Methods

Consider a MALDI-TOF IMS dataset *M* (∈ ℝ^*m×n*^), where *m* is the number of pixels/spectra, *n* is the number of *m/z* - bins recorded by the instrument, and *M*_*ij*_ is the ion intensity value associated with the *i*-th pixel and *j*-th *m/z* -bin. This dataset consists of real-valued ion intensities, where a row of *M* is a spectrum associated with a specific spatial location in the tissue and where a column of *M* is a particular *m/z* -bin considered across all pixels. The latter can be reconstructed into a so-called ion image, reporting the spatial distribution and abundance of a specific *m/z* -bin’s intensities. For some IMS experiments, the intensity values *M*_*ij*_ are clipped by the instrument (*e*.*g*., explicitly by acquisition-time data reduction or implicitly by the instrument’s limit-of-detection), effectively not reporting intensity values below a certain relative ion count *k*. Ion intensity clipping is sometimes expressly performed to induce a sparse regime^1^ on the recorded signals, often to save disk space. Regardless of the reason for clipping to occur, we want to explicitly deal with the missing values introduced by it. Therefore, we propose to model clipping as a function *f* : ℝ^*m×n*^→ ℝ^*m×n*^,

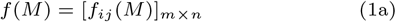

where *f*_*ij*_ (*M*) is defined for each entry (*i, j*) as:

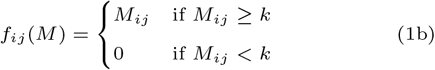

for *i* ∈ [1, *m*] and *j* ∈ [1, *n*]. The resulting (sparsified) dataset *f* (*M*) ∈ ℝ^*m×n*^ can be stored in a sparse data matrix format, a data structure that only explicitly stores non-zero values and their locations in the matrix, leaving zero values to be implicitly represented without consuming memory. Most post-acquisition data reduction methods ignore the non-linear operator, *f* (·), and treat the dataset *f* (*M*) as if it was *M* by assuming zero ion intensity for entries where *M*_*ij*_ *< k*. This is, for example, common in most peak picking algorithms, as they often require a full spectrum profile with an intensity value for each *m/z* -bin to determine where peaks are located. This rather naive approach may lead to information loss, assuming zero intensity where the abundance was low but not zero, and it can ultimately lead to biased feature subset selection, overemphasizing the importance of medium-to-high-abundant molecular species and underrepresenting or ignoring low-abundant species. In contrast, the methods presented here do not ignore the non-linear clipping operator, but rather seek to take it explicitly into account and avoid some of the assumptions listed above. Concretely, we propose to model the non-linear clipping function and describe it as a sampling operator. By applying a sampling operator 𝒫_Ω_(·) to *M*, essentially a relaxation for *f* (*M*), we acknowledge that there are missing values in *M* and avoid the assumption that those missing values are all zeroes when modeling. Yet, the sampling operator also avoids that those missing values (potentially) negatively impact the model. Specifically [Candes and Plan, 2010], 𝒫_Ω_ : ℝ^*m×n*^ → ℝ^*m×n*^ is defined as

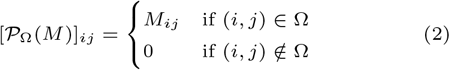

for *i* ∈ [1, *m*], *j* ∈ [1, *n*] and where Ω is the set of indices corresponding to the (known, reported) sampled values, as obtained from the instrument and after any acquisition-time data reduction. We denote (*i, j*) ∉ Ω as the set Ω_*c*_, making Ω and Ω_*c*_ complementary subsets of all entries in *M*. We can formulate the modeling of an IMS dataset *M* implicitly as a missing value problem, wherein a low-rank matrix approximation *X* of *M* is sought in the presence of missing data. The *M* -approximating matrix *X* can be considered an underlying model for the observed measurements in *M*, and the rank of *X* denotes the dimension of the subspace containing the approximating matrix. Low-rank matrices have several favourable mathematical properties in the context of underdetermined systems of equations and are oftentimes used to describe data from the smallest possible set of basis vectors, *i*.*e*., “the simplest representation for the given measurements”. We formulate the modeling of IMS data, *i*.*e*., the missing value problem, as an optimization problem that seeks to capture the IMS data using as little rank as possible, while concurrently being aware of the sampling operator and thus missing values:

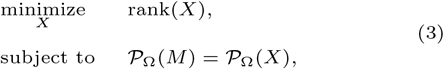

where *X* ∈ ℝ^*m×n*^ and 𝒫_Ω_(·) : ℝ^*m×n*^ → ℝ^*m×n*^ can be seen as an orthogonal projection projecting a matrix onto the space of ℝ^*m×n*^ matrices with support Ω. Since this optimization problem is non-convex (NP-hard), difficult to calculate, we instead solve a convex relaxation of the problem in Eq. 3 using the singular value thresholding (SVT) algorithm [Cai et al., 2010, Candes and Plan, 2010]:

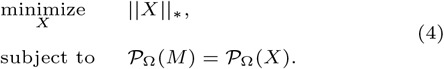

This program is shown to exactly recover the problem in Eq. 3 under specific conditions, *e*.*g*., incoherence of bases and sampling distribution [Candes and Recht, 2012]. As proving a condition’s validity is considered to be as hard as solving Eq. 3, the conditions cannot be verified. Thus, we will assume that conditions are met. Nevertheless, we will discuss the sampling distribution assumption in the Case Study section of this paper, as it is closely intertwined with the clipping mechanism during acquisition-time data reduction.

A strong advantage of the SVT is that during the optimization it utilizes the matrices in either a sparse or low-rank format, and it does not require a dense copy to be stored in memory. This is an extremely important aspect of this modeling effort, since many IMS datasets simply do not fit in memory when considered in a dense data format. A downside, however, is the singular value decomposition (SVD) at the center of each iteration of the optimization. Namely, the time complexity of the SVD, 𝒪(*mn*^2^), becomes a bottleneck at the scale of MALDI-TOF IMS datasets, mainly due to a number of BLAS level-2 operations at the heart of the SVD [Dongarra et al., 2018]. However, different solutions exist to reduce, *e*.*g*., [Halko et al., 2011]), or completely remove the SVD, *e*.*g*., [Zhou and Tao, 2011]. We opt for a divide-factor-conquer approach [Mackey et al., 2015], as it is theoretically well-studied and acts as a framework that we can adapt for a second method.

Besides the sparse-format-aware SVT approach and its SVD-related modification to calculate our low-rank approximation of IMS data, we also explore a second method that has a similar solving program as in Eq. 4, but it relaxes the equality into an inequality constraint:

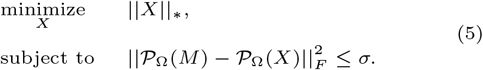

The latter enables the use of a fixed point continuation (FPC) algorithm for solving instead [Candes and Plan, 2010, Ma et al., 2011]. This second program accounts, in addition to missing values, for low-intensity dense noise (*e*.*g*., Gaussian noise) in the measurements, which is also inherently present in IMS data. The disadvantage of this algorithm is that it requires a dense-format matrix of similar size as *M* to be stored in memory (for the MALDI-TOF IMS dataset in this paper, this amounts to 1649.769 GB). As such, it is clear that whether SVT or FPC are the better choice for solving these optimization problems depends on the resources available and the needs of the subsequent analysis. Finally, note that both SVT and FPC’s practical implementations have a *δ* and *τ* parameter that arise as part of their solving algorithms. These hyperparameters require *a priori* setting (or optimization). We specify their setting for each experiment in Addendum A.4.

Furthermore, to deal with both SVD complexity and memory load, we make use of the divide-factor-conquer approach (DFC) [Mackey et al., 2015]. It consists of three steps and provides a framework that we can apply to both the SVT and FPC algorithms for obtaining an approximation 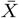of matrix *M* with completion for the complete dataset *M*. Its first step consists of dividing *M* into *t* different matrices *C*_*e*_, consisting of a subset of *M* ‘s columns, sampled uniformly at random:

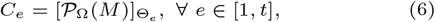

where Θ_*e*_ consists of a set of column indices of size 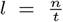 and Θ contains *t* such sets. We assume for ease that *l* is an integer. As such, we obtain *t* matrices *C*_*e*_ ∈ ℝ^*m×l*^. Note that each *C*_*e*_ also has a particular 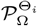 associated with it, namely the sampling associated to those columns in Θ_*i*_. In the second step, these subsampled matrices are factorized separately using the matrix completion methods, either SVT or FPC. Hence, *t* low-rank matrices 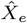 are obtained. The rank of the overall approximation 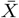 is calculated by taking the median of all matrix ranks of 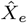. This rank is utilized in the final step, which consists of reconstructing the factored solutions 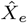 into a final approximate factorization 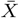. A standard Gaussian matrix *G* ∈ ℝ^*m×*(*k*+*p*)^ is constructed, with *p* as oversampling parameter, generally used to improve the reconstruction [Halko et al., 2011]. Next, a power iteration scheme is implemented as 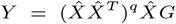, with *q* as the number of iterations and 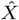 consists of re-ordering and stacking all low-rank approximations. Finally, the top *k* singular values of *Y* are obtained, *e*.*g*., by QR decomposition, to form *Q* ∈ ℝ^*m×k*^. The final solution 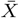 is then obtained by

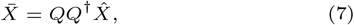

where † is representing the pseudo-inverse. The advantage of the divide-factor-conquer approach is that conditions that arise in matrix completion methods, as presented earlier, also guarantee strong estimation properties for divide-factor-conquer [Mackey et al., 2015]. The specifics of subsampling are provided in the Addendum A.4. We implemented the SVT and FPC algorithms as well as the divide-factor-conquer approach for both in an efficient Python object-oriented toolbox, with an eye towards saving memory where most needed and accelerating calculations where possible.

## Case Studies

In a first case study, we establish that an IMS dataset representation using a low-rank matrix factorization approach can outperform an equally small IMS dataset representation using traditional peak picking in a no-missing value case. We demonstrate this on Fourier-transform ion cyclotron resonance (FT-ICR) imaging mass spectrometry data (**Fig. 1**). In the second case study, we investigate the reconstruction error (*i*.*e*., on sampled/known values, ∈ Ω), the imputation error (*i*.*e*., on missing values, ∈ Ω_*c*_) and the global error (*i*.*e*., on all entries, ∈ (Ω ∪ Ω_*c*_)) on the same FT-ICR IMS dataset as in the first case study (**Fig. 1**). To mimic missing entries in the FT-ICR data, we implement two sampling schemes. The goal of the third case study is to evaluate the methodology, specifically the SVT and FPC algorithms with the divide-factor-conquer approach, directly on TOF IMS data that inherently includes missing values (**Fig. 2**). This dataset consists of 312, 249 *m/z* -bins for 1, 320, 876 spectra (1.65 TB in dense matrix format). The evaluation is both quantitative, using an error score and compression factor, and qualitative, with a focus on visualizing advantages and limitations that are relevant to analytical chemists. The data preprocessing can be found in Appendix A.3.

**Fig. 1.**
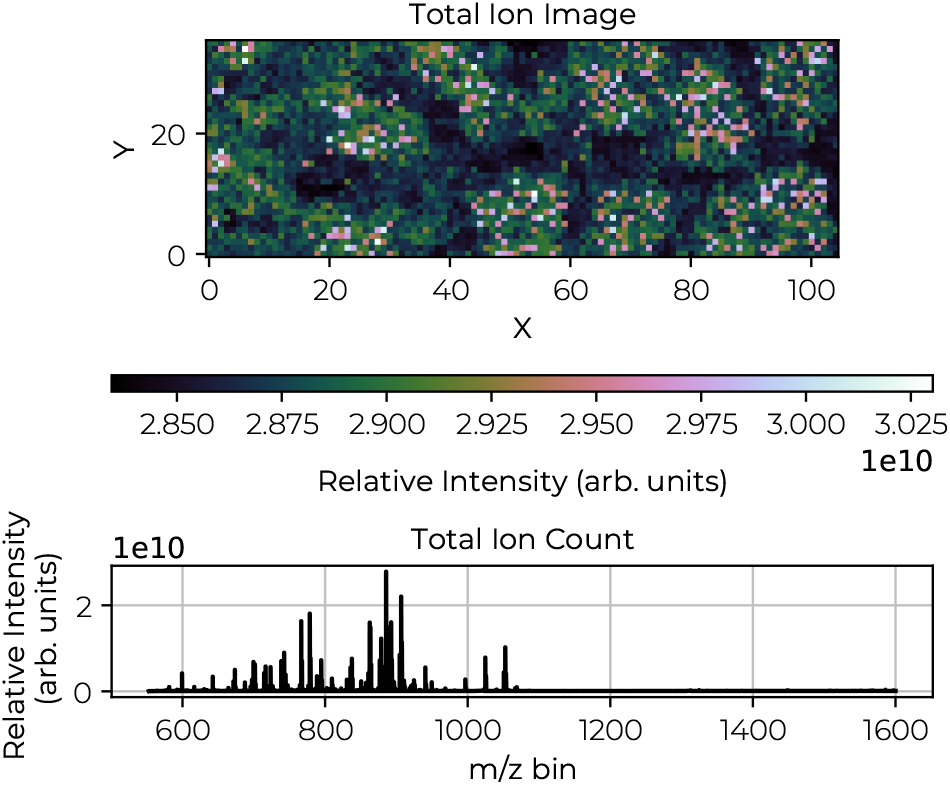
MALDI FT-ICR IMS measurement of human kidney tissue. The experiment was conducted using a 15T Bruker MALDI FT-ICR mass spectrometer (Bruker Daltonics, Billerica, MA, USA) with 50-*μ*m pixel size, covering the *m/z* range from 552 to 1,600 in negative ionization mode. For further sample preparation specifics, see Addendum A.1. The raw data were exported to a custom file format and normalized using 5-95%-TIC. The dataset contains 3, 780 spectra, each consisting of 1, 372, 421 *m/z* -bins. For further data preprocessing specifics, see Addendum A.3. The top panel shows the spatial distribution, represented as a total ion current image (*i*.*e*., the summation of the normalized spectral axis). The bottom panel displays the average mass spectrum.

**Fig. 2.**
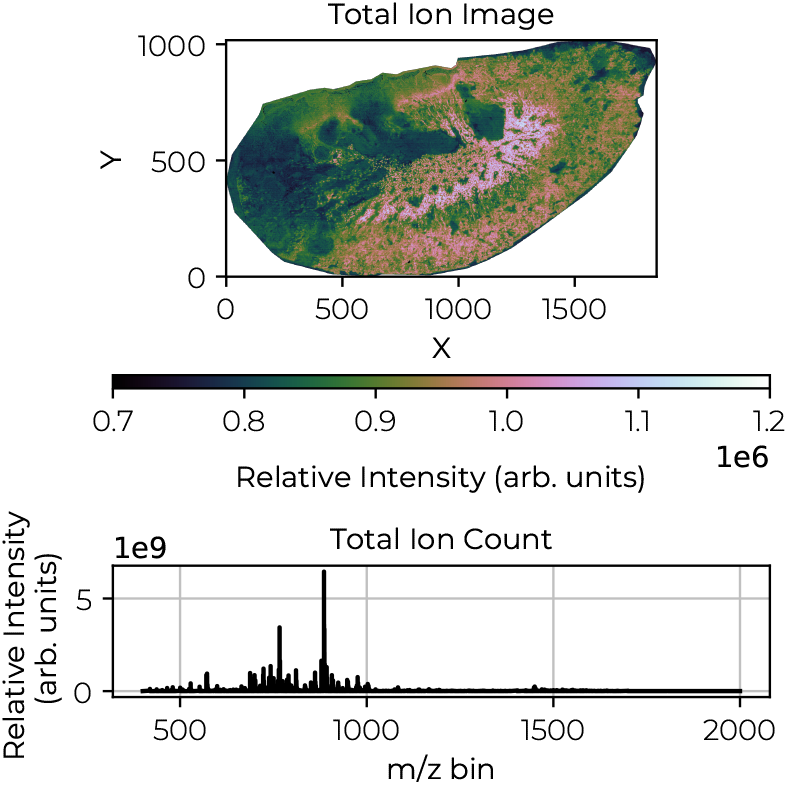
MALDI qTOF IMS measurement of *Staphylococcus aureus*-infected mouse kidney tissue. The infection-induced abscesses are visible as dark green areas in the total ion current image. The experiment was performed using a Bruker timsToF Flex mass spectrometer (Bruker Daltonics, Billerica, MA, USA) with 5-*μ*m pixel size, covering *m/z* range from 400 to 2,000 in negative ionization mode. For further sample preparation specifics, see Addendum A.2. The raw data were exported to a custom file format and normalized using 5-95%-TIC. The dataset contains 1, 320, 876 spectra, each consisting of 312, 249 *m/z* -bins. For further data preprocessing specifics, see Addendum A.3. The top panel shows the spatial distribution of the total ion current image. The bottom panel displays the average spectrum.

### Case Study 1: Low-Rank Matrix Factorization Outperforms Traditional Peak Picking

We first demonstrate that a low-rank matrix approximation can achieve a lower reconstruction/global error compared to peak picking in a no-missing value case, in addition to retaining full spectrum information. Having established this baseline, we can expand our problem setting with missing values in the second case study.

The best rank-*k* approximation with respect to the Frobenius norm (a measure we will use throughout this paper) is given by the truncated singular value decomposition (SVD) [Eckart and Young, 1936, Mirsky, 1960]. Since peak picking can be viewed as a form of (low-rank) matrix approximation by selecting specific columns from a dataset (with a column representing a selected peak), we can assert that peak picking is, at best, as effective as the truncated SVD. In **Table 1**, we observe a 39.1% difference in reconstruction error (which is equal to the global error in the absence of missing values) between the truncated SVD (factorization) and peak picking (100 peaks). If we compare the same reconstruction error based on a similar data footprint, this would still amount to a difference of 36.8%. While a reconstruction score can be a rather abstract form of capturing full spectrum information content, we highlight in **Fig. 3** a concrete difference between the raw IMS data, a low-rank representation (rank-100), and the conventional peak-picked representation. This example demonstrates that not only the overall ion intensity profile is preserved in the factorization representation, but a low-abundant peak at *m/z* 778.524 is effectively retained, where the peak-picked representation misses this peak entirely. Both the simple metric used in **Table 1** and the example of missing low-abundance peaks in **Fig. 3** illustrate that low-rank matrix factorization outperforms traditional peak picking when it comes to IMS dimensionality reduction. However, while SVD is a strong factorization method, it may not always be optimal, *e*.*g*., when dealing with missing values in the data.

**Table 1.**
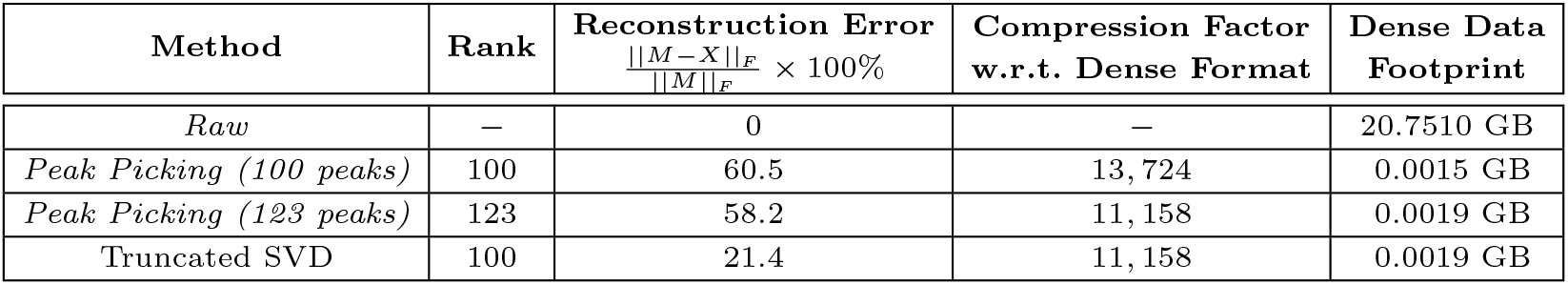
Truncated SVD and peak picking results. The reconstruction error is used throughout this paper as metric to measure how well (full) spectrum information is captured. A larger error implies that more information is lost. Hence, a low error is desired. However, note that an error of 0% is (probably) not desired as the data contains noise and it would be desirable to filter off this noise, leading to a (small) error. From this table, we observe that a factorization approach, the truncated SVD, leads to a substantial decrease in reconstruction score (up to 39.1%) compared to peak picking. The truncated SVD is carried out by truncating an SVD performed by the GESDD-routine. Peak picking is performed by matching (1) the rank and (2) the data footprint, *i*.*e*., MBs on disk. The raw data in dense matrix format has a storage footprint of 20.751 GB.

**Fig. 3.**
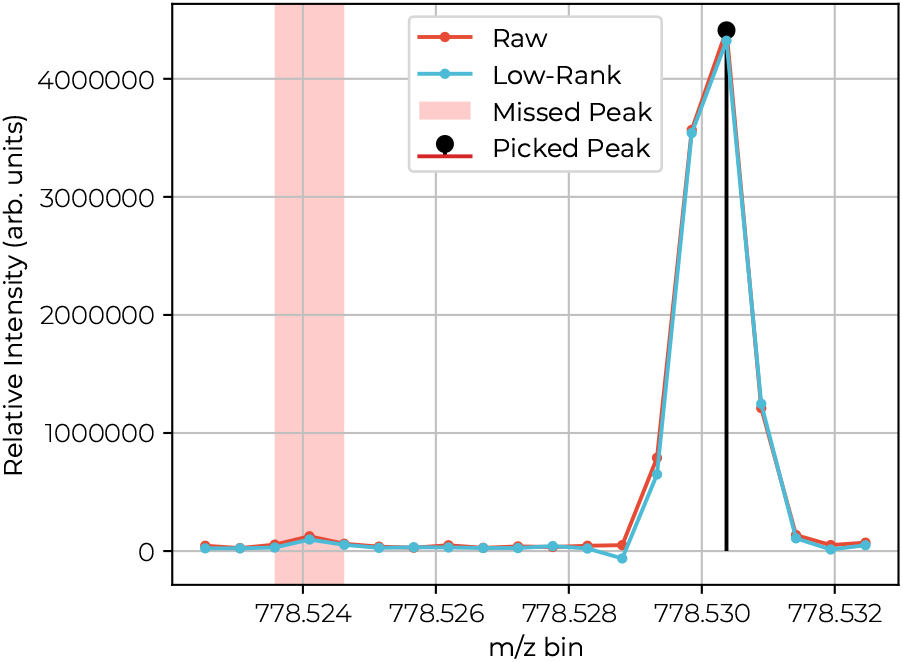
MALDI FT-ICR spectrum of a particular pixel showing the raw, low-rank and peak-picked data in a particular *m/z* -window of the mass spectrum. The area, highlighted in red, shows a low-abundant peak missed by the peak-picking representation, yet accurately captured and reconstructed by the low-rank factorization representation. This is achieved at identical compression ratios for both representations. If low-abundant peaks are of interested, factorization-based representations are probably better suited to reduce the dimensionality of IMS datasets.

### Case Study 2: Reconstruction and Imputation Quality when Dealing with Missing Values

Having established that a factorization approach is favourable over peak picking in a no-missing value situation, our factorization approach is now evaluated in a missing value situation. For this case study, we therefore implement two sampling schemes to mimic missing values in IMS data:

(*α*) Selects the top 8.9% of intensity values (to establish the in-sampling set Ω, *i*.*e*., the known values), with all other (lower) intensity values removed (making up the out-of-sampling set Ω_*c*_, *i*.*e*., missing values),

(*β*) Selects 8.9% of entries, not based on intensity but, uniformly at random (in-sampling set Ω, *i*.*e*., the known values), with all other values removed (out-of-sampling set Ω_*c*_, *i*.*e*., missing values).

The scheme *α* mimics commonly employed IMS acquisition-time data reduction, while the scheme *β* examines discrepancies related to incoherence conditions imposed by most matrix completion algorithms [Candes and Plan, 2010, Candes and Recht, 2012]. The 8.9% sampling rate was chosen for IMS fidelity to match the TOF IMS dataset sampling rate presented in the third case study. **Table 2** presents error scores for both the SVT and FPC algorithms (without the divide-factor-conquer approach) using sampling scheme *α* on the FT-ICR IMS dataset and reports:

**Table 2.**
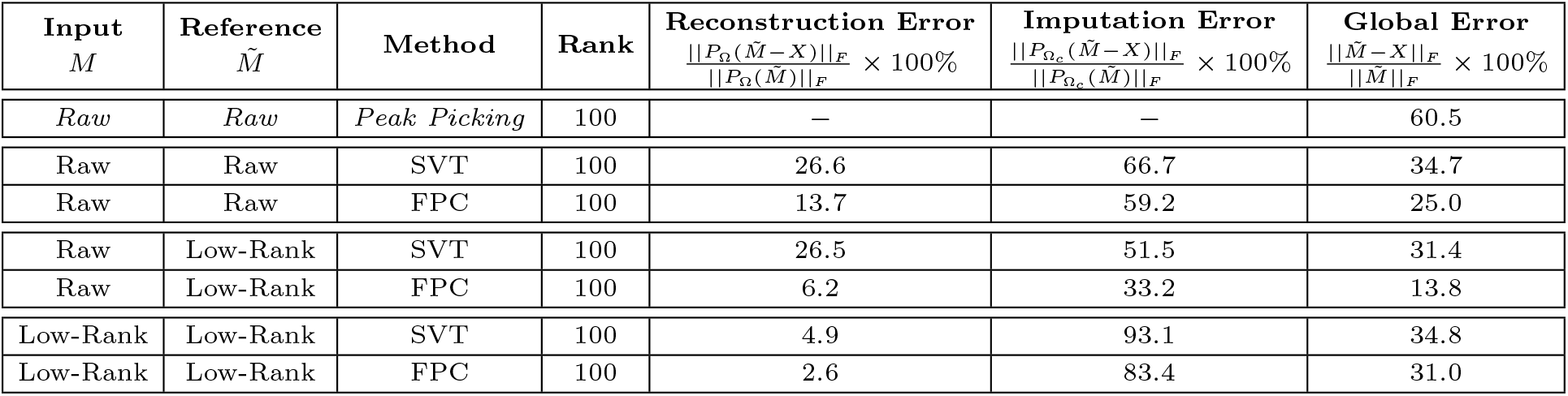
SVT and FPC results, with threshold sampling scheme *α*. A low reconstruction error is observed for all methods for both raw and low-rank input and references, comparable to the no-missing values case. The imputation error is more substantial. However, since these consists mostly of low intensity values (caused by the clipping operator) its impact is small on the global error. For SVT, we set parameters *δ* = 1 and *τ* = 10^*−*3^ and for FPC, we set *δ* = 1.4 and *τ* = 10^*−*3^ (see Addendum A.4). Peak picking is performed by picking the 100 highest peaks of the total ion current count of the raw data. For SVT with raw input data, we obtain a 171 rank solution, and for FPC, a 271 rank solution. For SVT with low-rank input data, we obtain a 131 rank solution, and for FPC, a 111 rank solution. We truncate all solutions to a rank of 100 for fair comparison.

| Input<br>$M$ | Reference<br>$\tilde{M}$ | Method | Rank | Reconstruction Error | Imputation Error | Global Error |
| --- | --- | --- | --- | --- | --- | --- |
| | | | | $\frac{\ P_{\Omega}(\tilde{M}-X)\ _F}{\ P_{\Omega}(\tilde{M})\ _F} \times 100\%$ | $\frac{\ P_{\Omega_c}(\tilde{M}-X)\ _F}{\ P_{\Omega_c}(\tilde{M})\ _F} \times 100\%$ | $\frac{\ \tilde{M}-X\ _F}{\ \tilde{M}\ _F} \times 100\%$ |
| Raw | Raw | Peak Picking | 100 | — | — | 60.5 |
| Raw | Raw | SVT | 100 | 26.6 | 66.7 | 34.7 |
| Raw | Raw | FPC | 100 | 13.7 | 59.2 | 25.0 |
| Raw | Low-Rank | SVT | 100 | 26.5 | 51.5 | 31.4 |
| Raw | Low-Rank | FPC | 100 | 6.2 | 33.2 | 13.8 |
| Low-Rank | Low-Rank | SVT | 100 | 4.9 | 93.1 | 34.8 |
| Low-Rank | Low-Rank | FPC | 100 | 2.6 | 83.4 | 31.0 |

**Table 3.**
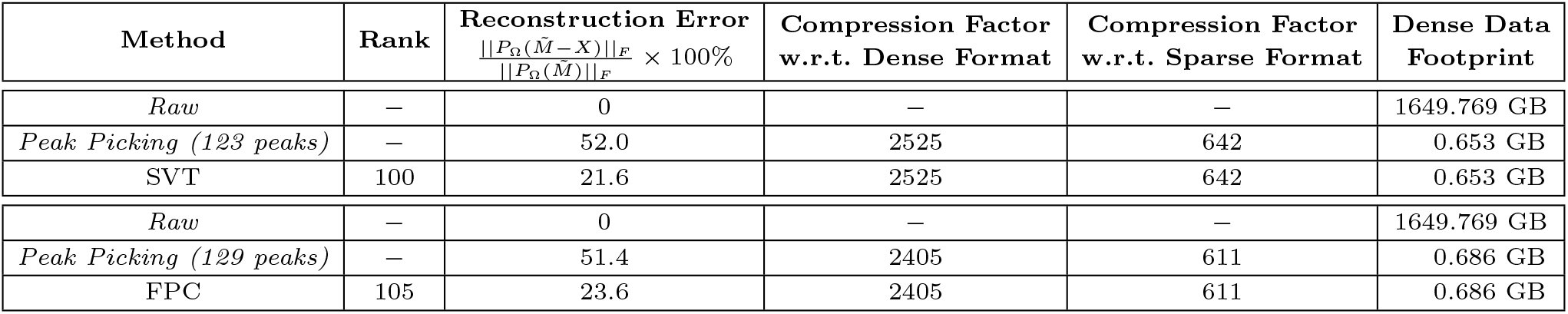
SVT and FPC, with randomized projection divide-factor-conquer approach. A substantial improvement for the reconstruction error is observed for both SVT and FPC in comparison to peak picking, while compression factors and data footprint are equal. For SVT, we set parameters *δ* = 1.7 and *τ* = .5 and for FPC, we set *δ* = 1 and *τ* = 1.5 × 10^*−*2^ (see Addendum A.4). Peak picking matches the data footprint, *i*.*e*., MBs on disk. The raw data in dense matrix format has a storage footprint of 1649.769 GB, and storing it in a sparse matrix format (*e*.*g*., compressed sparse column) amounts to 279.648 GB.

- Reconstruction error: modelling error for known entries.
- Imputation error: modelling error for missing values.
- Global error: modelling error for both known and missing entries.

Error scores were calculated with respect to both raw data and its low-rank approximation. Therefore, the input matrix is defined as the matrix used as input to our algorithms (Eq. 4, 5). The reference matrix is defined as the matrix used as reference in the error scores. We considered two types of input (*M*) and reference 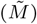 matrices:

- Raw: A dataset with missing values sampled directly from the raw data (thus including high-rank noise variation),
- Low-rank: A dataset with missing values sampled from a low-rank version of raw data, obtained through truncated singular value decomposition with rank 100.

Using the low-rank approximation as input (*M*) and reference 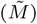 ensures that the low-rank conditions imposed by SVT and FPC are met, reducing unwanted (often noisy) variation from impacting the evaluation process.

Finally, note that the presented results stem from single experiments influenced by various factors (*e*.*g*., tissue type, sample preparation, detector type, raw data structure, noise levels, algorithmic parameters). Consequently, they are only evaluated relative to each other.

### Reconstruction, Imputation and Global Error

As shown in **Table 2**, both methods exhibit relatively low *reconstruction error* for non-missing values (between 2.6 and 26.6%) across all input and reference matrices. This is comparable/a slight improvement with respect to the results found in the first case study. Generally, FPC outperforms SVT on the raw input matrix, which is expected due to FPC’s ability to filter out small dense noise. On the other hand, both methods show only moderate performance on *imputation error* for missing values (between 33.2 and 93.1%) across all input and reference combinations, with FPC showing a slight advantage. This trend, along with similar results from the uniform sampling scheme *β* (**Table 4** in Appendix), suggests that while a low-rank factorization representation does a great job for capturing non-missing value entries, its performance as a predictor for missing values is limited and further investigation is needed to better understand the underlying causes.

**Table 4.** SVT and FPC, with random sampling scheme *β*. A moderate reconstruction error is observed for all methods for both raw and low-rank input and references. The imputation error is substantial. For sampling scheme *β*, the impact of the imputation error is larger on the global error. For SVT, we set parameters *δ* = 1 and *τ* = 10^*−*3^ and for FPC, we set *δ* = 1.4 and *τ* = 10^*−*3^ (see Addendum A.4). For SVT with raw input data we obtain a 171 rank solution and for FPC a 271 rank solution. For SVT with low-rank input data we obtain a 171 rank solution and for FPC a 271 rank solution. For the truncated SVD we used the lowest rank of the other experiments, *i*.*e*., 171. We truncate all solutions to a rank of 100 for fair comparison.

| Input<br>$M$ | Reference<br>$\tilde{M}$ | Method | Rank | Reconstruction Error | Imputation Error | Global Error |
| --- | --- | --- | --- | --- | --- | --- |
| | | | | $\frac{\ P_{\Omega}(\tilde{M}-X)\ _F}{\ P_{\Omega}(\tilde{M})\ _F} \times 100\%$ | $\frac{\ P_{\Omega_c}(\tilde{M}-X)\ _F}{\ P_{\Omega_c}(\tilde{M})\ _F} \times 100\%$ | $\frac{\ \tilde{M}-X\ _F}{\ \tilde{M}\ _F} \times 100\%$ |
| Raw | Raw | SVT | 100 | 50.0 | 92.6 | 89.6 |
| Raw | Raw | FPC | 100 | 29.3 | 84.9 | 81.6 |
| Raw | Low-Rank | SVT | 100 | 48.3 | 92.2 | 89.2 |
| Raw | Low-Rank | FPC | 100 | 23.5 | 84.2 | 80.7 |
| Low-Rank | Low-Rank | SVT | 100 | 67.9 | 92.4 | 90.5 |
| Low-Rank | Low-Rank | FPC | 100 | 22.8 | 84.4 | 80.9 |

Interestingly, contrary to expectations, the uniform sampling scheme *β* does not outperform the threshold-based sampling *α*, despite its closer alignment with incoherence conditions. This highlights the need to carefully consider the implications of different sampling strategies. Threshold sampling scheme *α* removes low-intensity entries, primarily associated with noise. Hence, it requires the imputation of noisy features by a low-rank model. However, this scheme generally fails to satisfy incoherence conditions, leading to poor imputation error in general. The poor performance could, for example, be caused by the spatial correlation of low-abundance values, which is particularly evident in specific *m/z* -bins and distinct (positive) spatial areas across the tissue. As illustrated in **Fig. 4**, under an intensity-magnitude driven sampling scheme *α*, a high-intensity *m/z* -bin at 885.571 is missing only a few values (white entries), while a low-intensity *m/z* -bin at 756.254 can be missing many values.

**Fig. 4.**
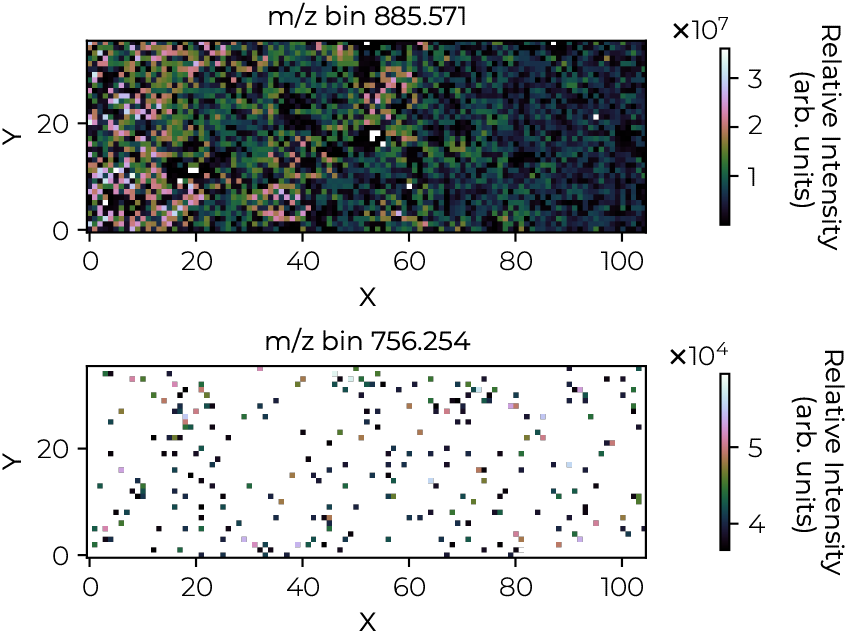
MALDI FT-ICR ion images from a single *m/z* -bin of the low-rank input matrix with threshold sampling scheme *α*. The top image depicts an *m/z* -bin with high intensity and thus, with scheme *α*, a low number of missing values. The bottom image depicts an *m/z* -bin with low intensity and thus, with scheme *α*, a high number of missing values.

In contrast, a uniform sampling scheme *β* is expected to perform better because it more closely aligns with incoherence conditions and treats all ion species the same. However, we observe worse reconstruction and imputation scores compared to scheme *α*. This is probably related to (1) the low 8.9% sampling rate (*i*.*e*., 91.1% of all intensity entries are missing in this dataset and there is relatively little signal to model with), and (2) the predominant number of low signal-to-noise *m/z* - bins in the raw data. Consequently, uniform sampling leads to a significant loss in high-valued, “informative” entries. We expect that the imputation error for sampling procedure *α* and both reconstruction and imputation error for sampling procedure *β* could benefit from advanced feature scaling. Additionally, introducing chemical noise (*e*.*g*., speckle noise) into our data before applying the sampling scheme could cause high-intensity values to become missing, albeit at a lower rate, which may further reduce the imputation error.

Nevertheless, both methods achieve a *global error* that is 25.7 to 46.7% lower than that of peak picking, which has a global error of 60.5%, representing a substantial improvement in terms of full spectrum information, even in the presence of missing values. Moreover, note that the imputation error does not significantly influence the global error score since low-intensity features contribute less to the global error—due to the properties of the Frobenius norm and because the sampling scheme *α* retains high-intensity values. Overall, this result implies that full spectrum information is better captured by the proposed low-rank factorization methodology than by peak picking, both when missing values are present as well as when they are absent.

### Case Study 3: Advantages and Disadvantages of Low-Rank Matrix Completion for Missing Value TOF IMS Data

In this case study, we forgo synthetically generated missing values, for which the ground truth is known, and apply our approach on IMS data with intrinsic missing values.

#### Reconstruction Error and Compression Factor

The performance metrics for this third case study, including the reconstruction error and compression factors, are summarized in **Table 3**. They highlight that both SVT and FPC exhibit comparable performance. While a reconstruction error of approximately 20% might seem substantial at first, it actually represents a 30% reduction in information loss compared to traditional peak picking, all while maintaining the same data footprint and enabling full profile analysis in downstream workflows. Furthermore, all methods demonstrate high compression factors, with the IMS representation’s footprint being approximately 2500 times smaller compared to a dense matrix format and 600 times smaller than a sparse matrix format, comparable to those achieved by peak picking.

#### Spectral Error Distribution and Biological Interpretation

We further investigate the distribution of reconstruction errors for individual spectra, referred to as the spectral error score (**Fig. 5.a** and **Fig. 5.c**). This score is calculated both for

**Fig. 5.**
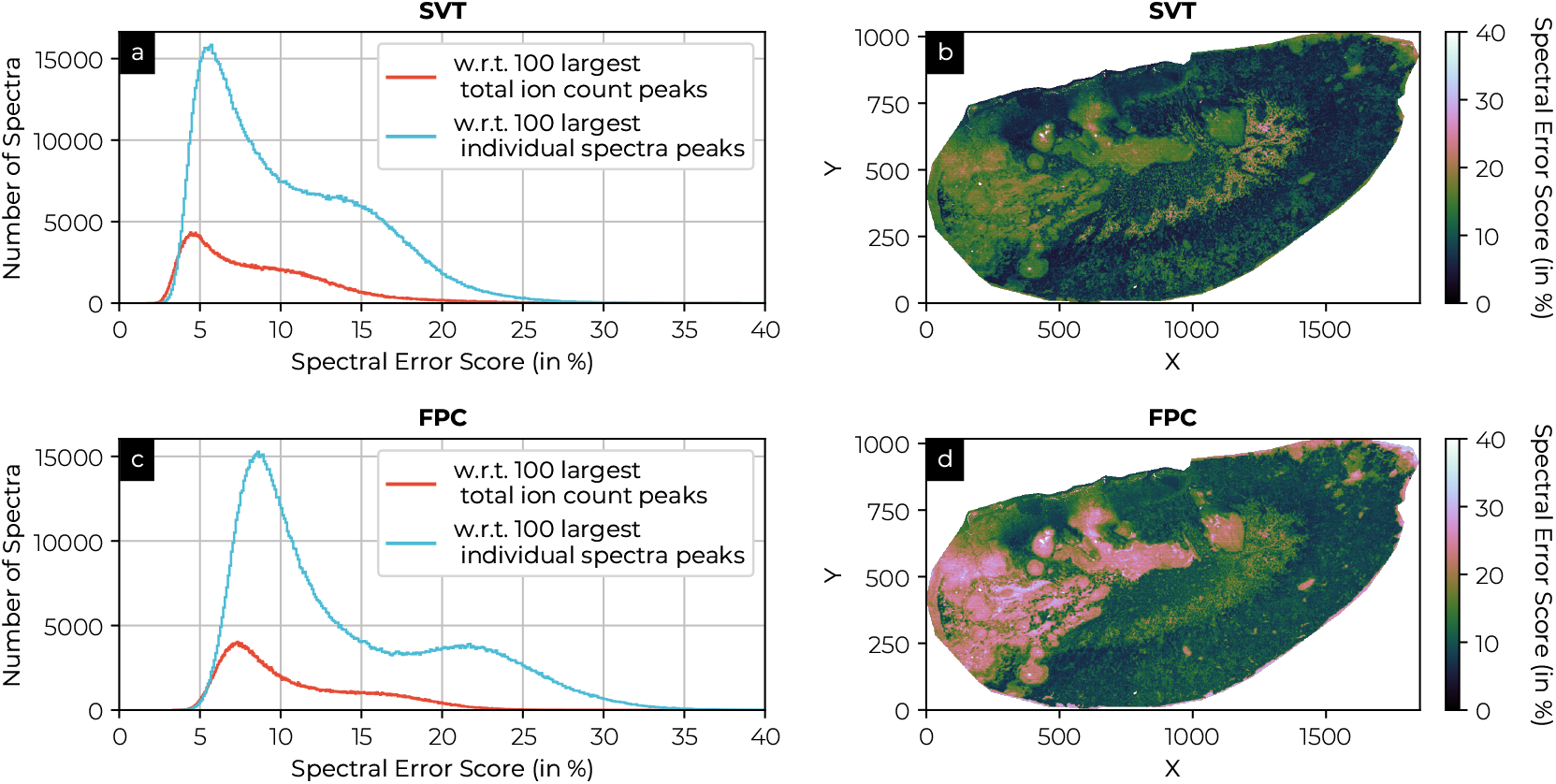
Spectral error score, *i*.*e*.,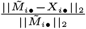, reports on the error of individual spectra. The distribution of the spectral error score is given (left) for both SVT and FPC methods with respect to the 100 largest total ion current count peaks and with respect to the 100 largest individual spectral peaks. The spatial distribution for the spectral error score with respect to the 100 largest individual spectra peaks is also depicted for the SVT and FPC (right).

(a) the 100 *m/z* -bins with the largest total ion current count across the dataset; and
(b) the largest 100 *m/z* -bins per spectrum, *i*.*e*., the top peaks in each individual spectrum.

The distributions of spectral error scores reveal patterns that correlate with biology for both SVT and FPC methods under both scoring criteria (*a* and *b*). Interestingly, the spectral error is slightly lower for the largest individual spectrum peaks (*b*), as the top dataset-wide peaks (*a*) might not be present in every spectrum. The error distributions appear to be a mixture of two Gaussian-like distributions with different means and standard deviations. Spatial reconstruction of these distributions (**Fig. 5.b** and **Fig. 5.d**) reveal distinct tissue regions that may correlate with the total ion current count (**Fig. 2**). Moreover, no clear relationship is observed between these distributions and the number of non-zero values per spectrum (**Fig. 9** in the Appendix). This suggests significant heterogeneity in molecular distributions within the tissue, rather than issues related to incoherence, might be influencing the reconstruction quality, especially in *Staphylococcus aureus*-infected regions.

#### Methodological Effects on Reconstructed Ion Images and Spectra

Although high-intensity ion images and peaks are recovered well (see reconstruction error and spectral error score), distortions may occur in the reconstructed spectra and individual ion images of very low intensity (**Fig. 6**). Commonly observed distortions included (1) small peak shifts, *i*.*e*., shifting of peak distribution along the *m/z* axis, (2) peak widening, *i*.*e*., smearing peaks over larger *m/z* ranges, and (3) peak prediction, *i*.*e*., imputation of peaks not present in the raw spectrum, but predicted on the basis of dataset-wide observed patterns. It should be noted that these effects are not necessarily incorrect, that they may arise from genuine corrections for small non-linear misalignments due to instrumentation or noise, or from other instrumental artifacts.

**Fig. 6.**
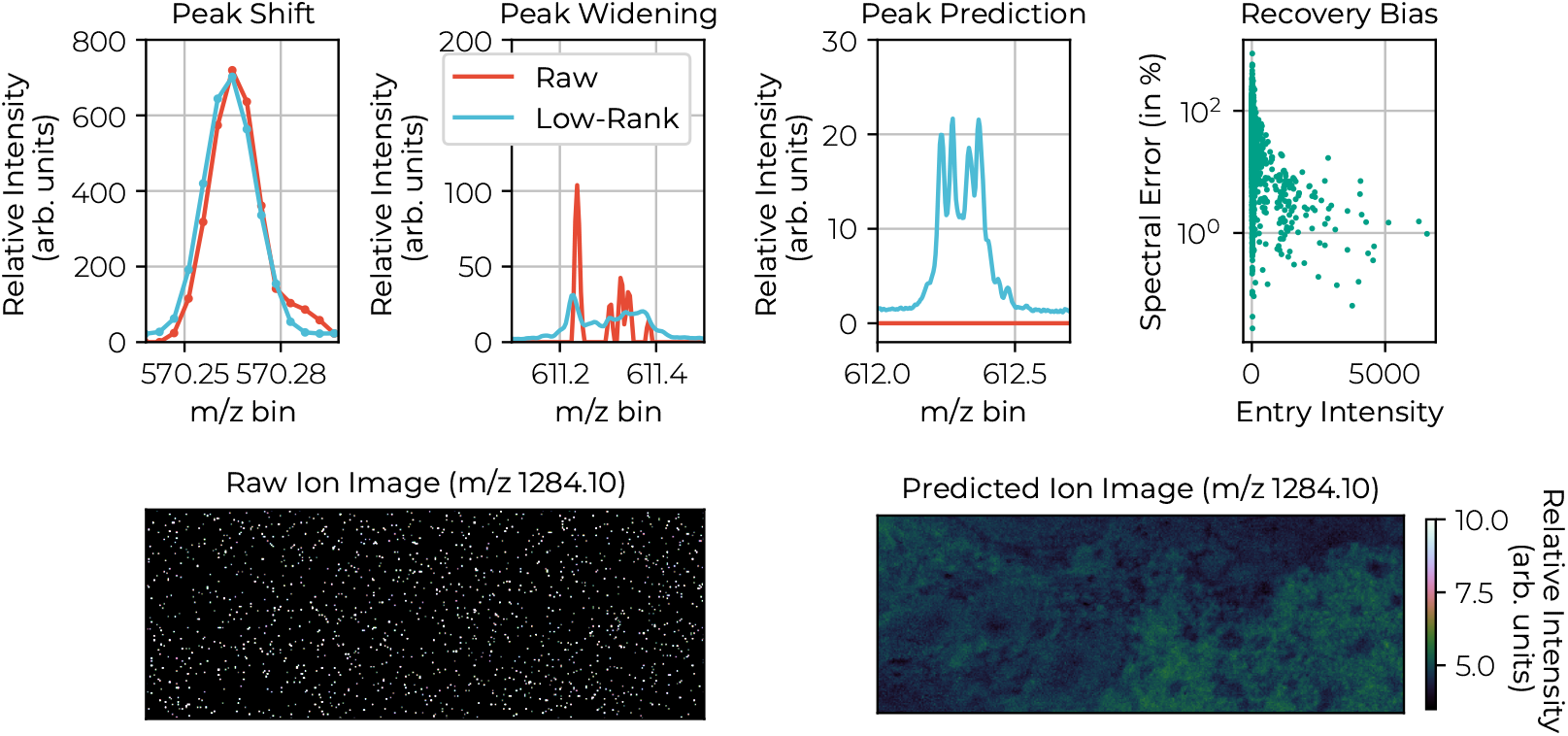
The first row depicts four potential effects of the low-rank approximation of a complete dataset on individual spectra, namely peak shifting (*i*.*e*., small shifts of the peak center), peak widening, peak prediction (potential introduction of low-intensity peaks in line with patterns observed in the rest of the dataset), and bias in the recovery error for large peaks (better recovery/representation of large peaks). In the second row, an extreme example is given of an individual ion image at *m/z* 1284.10, both raw (left) and recovered (right). The predicted image depicts a biological scene, even though the raw image contains barely any data points (very sparse). The missing values in the image are imputed, based on the available information from the rest of the dataset. This could lead to a substantial imputation error for those ion species. These effects are a price we pay for retaining full spectrum information in our dimensionality reduction methods, and they affect low-abundant and high-missing pixel species first. As such, we robustly keep the full spectrum profile intact for higher intensity peaks, in contrast to selective peak picking. At the same time, these recognized shortcomings can be taken into account for future model improvements.

Notably, these distortions are more prominent in low-intensity peaks, which is consistent with the optimization process focused on minimizing the Frobenius norm. This introduces (4) a recovery bias that favours better reconstruction of high-intensity peaks, as also observed in this TOF IMS dataset. From a spatial perspective, caution is warranted when interpreting very sparse ion images as biologically meaningful. For example, the predicted ion image of *m/z* 1284.10, is based on only very few measurements (see raw ion image of *m/z* 1284.10). These low-abundant species-centric effects, whether desired or undesired, can potentially be mitigated in the future through advanced feature scaling and an improved model. They should also always be considered within the context of peak picking approaches, which often leave no record of low-abundant species to begin with.

#### Preservation of Near-Isobaric Species

Near-isobaric species, molecular species with nearly identical mass-to-charge ratios but different chemical compositions, pose significant challenges for accurate peak detection. These species are often overlooked in peak picking due to their low intensity relative to dominant peaks, or may be incorrectly integrated as a single species. In **Fig. 7**, we present an example of such a near-isobaric species, at approximately *m/z* 725.53 (blue area), located close to a dominant species at *m/z* 725.51 (orange area). Due to their proximity, near-isobaric species are frequently neglected. Integrating the orange and blue areas separately, reveals different spatial molecular distributions, indicating that these *m/z* -ranges correspond to distinct molecular species. When applying peak picking, the best-case scenario consists of integrating the orange area and neglecting the blue area, potentially missing the near-isobaric species. In the worst-case scenario, both areas are integrated as one, and the dominant peak’s intensity overshadows that of the near-isobaric species, leading to the loss of unique spatial information. In both cases, the unique near-isobaric information is lost. However, our methods successfully preserve this information by approximating the full spectrum without requiring prior specification of their positions on the *m/z* -axis.

**Fig. 7.**
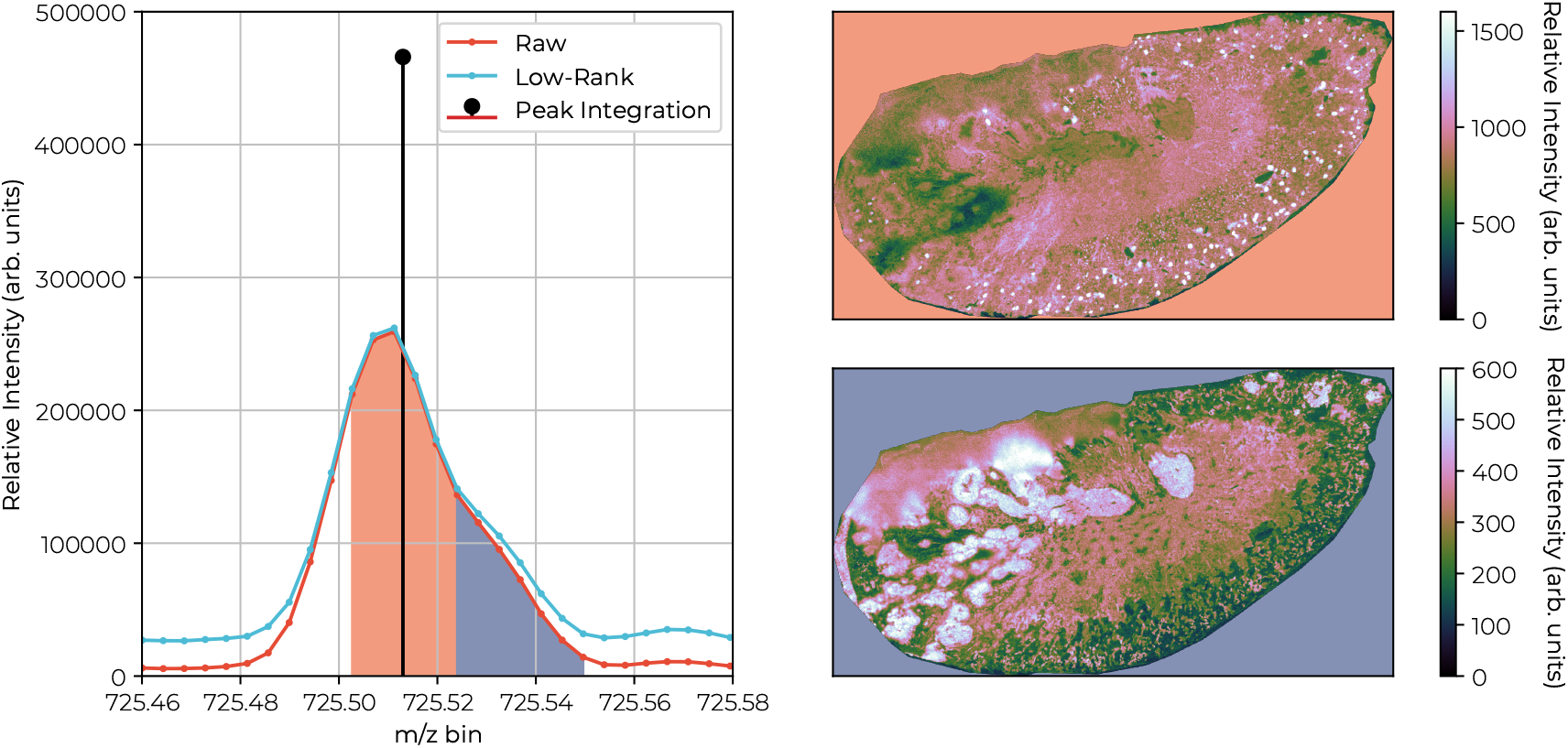
A particular spectrum is examined (left) in the range between *m/z* 725.46 and 725.58 from the perspective of both the low-rank approximation and raw data, along with a peak integrated version. The highlighted (orange and blue) areas under the curve are integrated and spatially depicted (right). We observe that the shoulder (blue) differs spatially from the peak (orange). With peak integration, relative information of the shoulder (blue) is lost, due to its weak signal.

This ensures that near-isobaric species are more accurately captured and represented, maintaining the unique spatial and molecular information they provide. Preserving near-isobaric species is an important factor in improving analysis specificity. Specificity on an instrumental and/or (bio)chemical level is an important driver in IMS [Spraggins et al., 2019], being able to maintain it in the analysis is thus of utter importance.

#### Retention of Lower-Intensity Ion Species and Bias Mitigation

Our approach effectively mitigates bias by retaining more lower-intensity ion species that are commonly disregarded by peak picking, especially when only the largest peaks are retained. We identified several *m/z* -bins representing peaks corresponding to biologically relevant lipids and adducts, which were preserved in our analysis despite their low intensity (**Fig. 8**). The specific *m/z* values include:

**Fig. 8.**
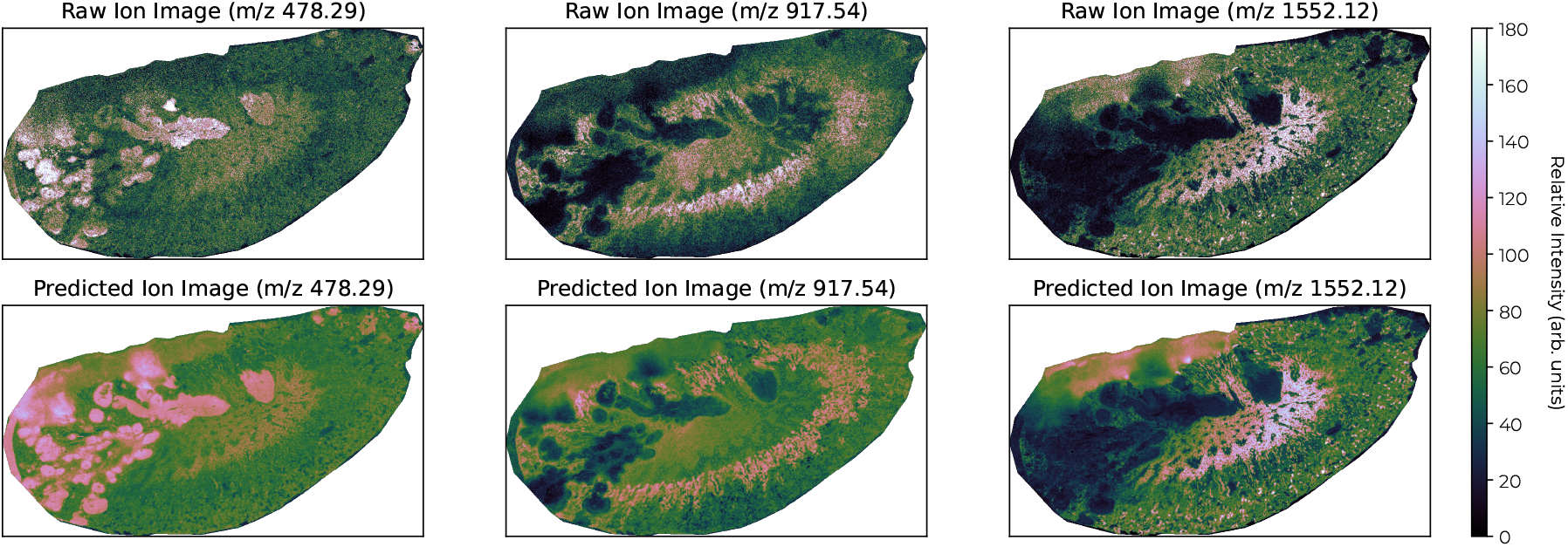
In the first row, the three columns depict different ion images as found in the raw data, namely *m/z* 479.29, *m/z* 917.54 and *m/z* 1552.12. These ion species are detected in more than 80% of the IMS pixels, they are not isotopes, and their distributions suggest a biology-driven distribution. Nevertheless, they would be disregarded in a simple peak picking procedure, if *e*.*g*., only the 1000 largest peaks would be retained. Our approach (see second row) retains these ion species, albeit approximately.

**Fig. 9.**
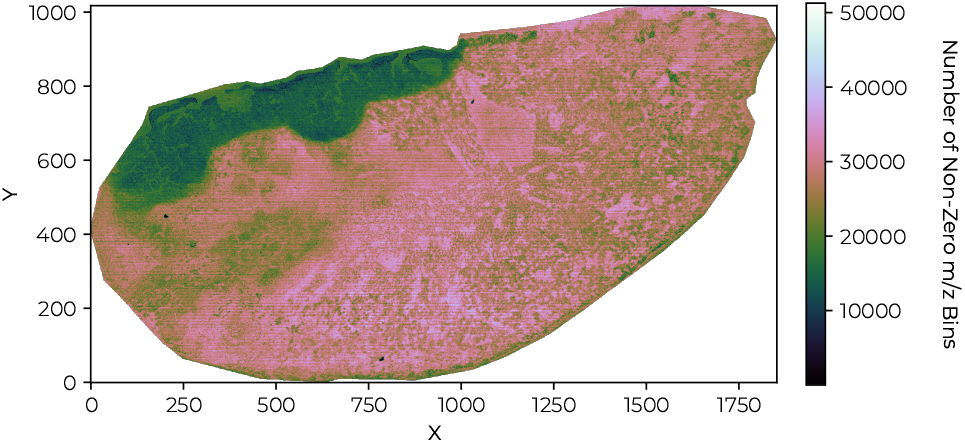
Spatial distribution of the number of non-zero values per spectrum. Dark regions correspond to spectra with few values (after clipping), while bright regions reflect spectra with more measured values. Note that most spectra only contain 2 *−* 30% of measured values. Some spatial regions contain more zeroes, *i*.*e*., fewer features are captured.

- The lipid LPE 18:1 (confirmed with LC-MS^2^) at *m/z* 478.29,
- A 4-(dimethylamino)cinnamic acid (DMACA) adduct^3^ of PE O-(36:3) at *m/z* 917.54,
- [CL(77:2)+Na-2H]-at *m/z* 1552.12.

These *m/z* -bins are not isotopic peaks and thus provide unique information about species abundant in different spatial regions in the tissue. These examples are only a few from a large group of peaks (1000+) that are preserved for this TOF IMS dataset. Retaining low signal-to-noise ratio signals enhances the detection of species in downstream analyses, thereby reducing confirmation bias by reporting on nearly all instrument-detected peaks. Retention of lower-intensity ion species in the computational representation and analysis is an important factor in maintaining sensitivity throughout the chain from sample preparation to instrument to computational analysis to biological insight.

## Conclusions

This paper explored the application of matrix factorization algorithms on IMS data, focusing on the goal of dimensionality and data footprint reduction, addressing the issue of missing values, and evaluating both quantitative and qualitative outcomes. For a no-missing value case, a low-rank factorization based representation of IMS data improved the reconstruction error by 39.1% over peak picking, while concurrently maintaining a full spectrum profile for all spectra in the dataset. In the missing value case, we achieved low reconstruction errors for both SVT and FPC based approaches, comparable to the no-missing case. We also highlighted the persistent challenge of reducing imputation errors, which could potentially be mitigated through advanced feature scaling that accounts for the specific characteristics of IMS data and an improved data model. For the missing value case, we demonstrated a substantial reduction in full spectrum information loss (global error) up to 40% compared to traditional peak picking methods, while achieving compression factors similar to peak picking. Our experiments revealed that matrix completion algorithms offer significant advantages in maintaining sensitivity by preserving lower-signal-to-noise ratio signals and mitigating selection bias. At the same time, we demonstrated preservation of specificity by preservation of near-isobaric species in the analysis through our full profile approach. Both are expected to impact downstream analysis by providing a richer and completer reduced representation of IMS data, while also providing dimensionality reduction capabilities comparable to traditional peak picking. The importance of this research lies in the introduction of a framework for IMS data reduction by factorization in an early stage and with awareness of missing values. This framework enables high compression rates, up to 2500-fold compared to dense matrix storage formats and up to 600-fold compared to sparse matrix storage formats, while preserving substantially more full profile information than peak picking. We emphasize the importance of utilizing full spectra in downstream analysis to avoid premature or biased information loss, as often occurs with peak integration or peak picking. However, our methods also have limitations, such as peak shifting and widening, low-intensity peak prediction, and the prediction of very sparse ion images.

Looking forward, future work could focus on exploring on-the-fly low-rank approximation schemes that can be employed during data acquisition to enhance accuracy and reduce computational burden. Additionally, it will be important to incorporate considerations for non-negativity, measurement sparsity, and uncertainty. Addressing issues related to peak shifting, widening, and normalization also emerges as a critical area for further research.

In conclusion, our study shows that low-rank factorization based representations of IMS data can substantially advance the field by reducing full spectrum information loss by 30 to 40% compared to traditional peak picking methods. This work highlights the potential of matrix factorization and, in particular, completion algorithms for avoiding premature feature selection and for lifting IMS data analysis to the full profile level.

## Appendix

### Sample Preparation: Human Kidney Tissue (FT-ICR IMS)

Human kidney tissue was surgically removed during a full nephrectomy, and remnant tissue was processed for research purposes by the Cooperative Human Tissue Network (CHTN) at Vanderbilt University Medical Center. Human biospecimens were collected in compliance with the CHTN protocols, institutional IRB policies, and the National Cancer Institute’s best practices for procurement of remnant surgical research material. The tissue was flash-frozen over an isopentane-dry ice slurry and embedded in carboxymethylcellulose. Tissue sections were cryosectioned with a thickness of 10 *μ*m and thaw-mounted onto indium tin-oxide-coated glass slides (Delta Technologies, Loveland, CO, USA). 1,5-Diaminonaphthalene (DAN) was applied to the tissue surface using a TM Sprayer M3 (HTX Technologies, Chapel Hill, NC, USA). The sample was imaged (50 *μ*m pitch) directly after matrix application with a 15T MALDI Fourier-transform ion cyclotron resonance (FT-ICR) mass spectrometer (Solarix, Bruker Daltonics, Billerica, MD, USA). Briefly, data were generated from m/z 300-2000 with a 4M file size in negative ion mode.

### Sample Preparation: Mouse Kidney Tissue (qTOF IMS)

C57BL6/J mice were retro-orbitally infected with *S. aureus* Newman and sacrificed humanely five days post-infection. Kidneys were flash frozen on an isopentane/dry ice slurry and embedded in 2.6% carboxymethylcellulose. 5 *μ*m thick sections were collected using a Leica Biosystems CM3050S cryostat and thaw-mounted onto indium tin oxide coated glass slides. Sections were washed with cold 150 mM ammonium formate for 45 seconds for three total washes. 5 mg of 4-(dimethylamino)cinnamic acid matrix was applied using an in-house sublimation device. MALDI IMS data were collected in negative ion mode from m/z 400-2000 using a Bruker MALDI timsTOF fleX platform with a 5 *μ*m step size, 25% laser power, and 25 shots per pixel.

### Data Preprocessing

The TOF dataset was *m/z* -aligned with a custom program [Migas, 2024, Monchamp et al., 2007, Farrow et al., 2022]. For all case studies, we used a 5-95% TIC pixel normalization method, *i*.*e*., scaling each row by the sum of its entries between 5%-percentile and 95%-percentile [Migas, 2024, Monchamp et al., 2007, Farrow et al., 2022]. We did not statistically normalize (*i*.*e*., mean subtraction and scaling) the features, *i*.*e*., *m/z* -bins (columns), as we observed that it hindered low-rank recovery. As the majority of measured *m/z* -bins consisted of low signal-to-noise measurements, sub-noise features or noise, statistically normalizing the *m/z* -bins boosts the singular values associated to the noise subspace, while dwarfing the singular values associated to the signal subspace, associated to high intensity *m/z* -bins. Exploring more advanced normalization schemes was considered out-of-scope for this paper. We performed all calculations on a Dell Precision 7920 with 56 cores at 2.7 GHz and 1.5 TB of memory with two NVIDIA A6000 GPUs with NVLink Bridge.

### Parameter Setting

For SVT, convergence for the completion problem is guaranteed if 0 *< δ <* 2 [Cai et al., 2010]. However, it was also noted that this choice can be too conservative, and the convergence slow [Cai et al., 2010]. For our datasets, we found that setting the parameter to *δ >* 2 breaks the convergence and that values of *δ <* 1 lead to slow convergence. Hence, we manually tuned the values per experiment between 1 *< δ <* 1.7. For FPC, different suggestions are made for setting *δ* [Candes and Plan, 2010, Ma et al., 2011]. We found by manually tuning that convergence was ensured for our datasets settings values, similar to SVT, between 1 *< δ <* 2.

Although different suggestions for SVT’s and FPC’s *τ* exist (note that *τ* is described by *μ* for FPC [Candes and Plan, 2010]), we defined *τ* relative to max (*m, n*). By trial-and-error, we found values between 10^*−*1^ and 10^*−*3^ perform adequate to obtain low-rank solutions. We did not observe large deviations in recovery when setting the parameters in those ranges. A parameter sensitivity analysis is, however, advisable to further tune the outcomes.

For the divide, factor and conquer approach, the raw data matrix *M* ∈ ℝ^312,249*×*1,372,421^ was divided into 3, 303 subsampled matrices, *C*_*i*_ ∈ ℝ^312,249*×*400^, where *i* ∈ [1, 3303]. Subsampling was performed along the spatial axis, which outperformed spectral axis subsampling due to the presence of many noise-dominant *m/z* -bins. Spectral subsampling risked aggregating noise features into the same subsampled matrices, leading to faulty results. Sampling sizes of 300 − 500 spectra were heuristically found to ensure good recovery, *i*.*e*., fulfilling the low-rank constraint while allowing for efficient computation, completing within a 12-hour runtime.

## Author contributions statement

## Acknowledgments

Research supported by the National Institutes of Health (NIH)’s Common Fund, National Institute Of Diabetes And Digestive And Kidney Diseases (NIDDK), and the Office Of The Director (OD) under Award Numbers U54DK120058, U54DK134302, and U01DK133766, by NIH’s Common Fund, National Eye Institute, and the Office Of The Director (OD) under Award Number U54EY032442, by NIH’s National Institute Of Allergy And Infectious Diseases (NIAID) under Award Numbers R01AI138581 and R01AI145992, by NIH’s National Institute On Aging (NIA) under Award Number R01AG078803, and by grant numbers 2021240339 and 2022309518 from the Chan Zuckerberg Initiative DAF, an advised fund of Silicon Valley Community Foundation

## Footnotes

1 In this context, sparsity refers to the number of non-zero values in measurements, *i*.*e*., high sparsity implies many zero values. A sparse regime implies that measurements contain many zero values.

2 Liquid chromatography–mass spectrometry

3 DMACA is a solution sprayed onto tissue during wet-lab preprocessing

